# Categorical signaling of the strongest stimulus by an inhibitory midbrain nucleus

**DOI:** 10.1101/2019.12.19.883702

**Authors:** Hannah M. Schryver, Malgorzata Straka, Shreesh P. Mysore

## Abstract

The isthmi pars magnocellularis (Imc), a group of inhibitory neurons in the vertebrate midbrain tegmentum, orchestrates stimulus competition and spatial selection in the optic tectum (OT). Here, we investigate the properties of relative-strength dependent competitive interactions within the barn owl Imc. Imc neurons exhibit switch-like as well as gradual response profiles as a function of relative stimulus strength, do so for competing stimuli both within and across sensory modalities, and signal the strongest stimulus in a dynamically flexible manner. Notably, Imc signals the strongest stimulus more categorically (with greater precision), and earlier than the OT. Paired recordings at spatially aligned Imc and OT sites reveal that although some properties of stimulus competition are correlated, others are set independently. Our results demonstrate that the Imc is itself an active site of competition, and may be the first site in the midbrain selection network at which stimulus competition is resolved.

## INTRODUCTION

For animals operating within complex environments, the ability to select the location of the highest priority stimulus is vital for adaptive behaviors. Stimulus priority is the combination of the physical salience of a stimulus as well as its behavioral relevance, and is computed at several sites in the brain [1]. Among these, the midbrain selection network has been studied for its important role in the control of stimulus selection for spatial attention [2, 3]. It consists of the superior colliculus (SC, in mammals, or optic tectum, OT, in non-mammals), which encodes a topographic map of multisensory and motor space, as well as a spatial map of stimulus priority [2, 3], and satellite brain areas in the midbrain tegmentum that are interconnected to the SC/OT [4–8].

Work in non-human primates has demonstrated that the intermediate-deep layers of the SC (SCid) are required for the selection of the target of spatial attention amidst distractors [9, 10]. In parallel, work in the avian midbrain has demonstrated that neurons in the OTid signal the highest priority stimulus categorically: they respond with a high firing rate when the stimulus inside their receptive field (RF) is of highest priority, but are suppressed abruptly to a low firing rate when it is no longer the highest priority [11–13]. Suppression of the responses of SCid/OTid neurons by competing stimuli has been reported in several vertebrate species [10, 14–21]. The source of such long-range competitive inhibition that underlies OTid’s categorical signaling has been identified to be a group of inhibitory neurons in the vertebrate midbrain tegmentum, called nucleus isthmi pars magnocellularis (Imc; [4–6]. Specifically, it has been shown in birds that inactivation of the Imc abolishes all competitive interactions in the OTid [22, 23], as well as in the cholinergic Ipc (isthmi pars parvocellularis; [7, 8, 24], another key area in the midbrain selection network, which serves as a point-to-point amplifier of activity across the OTid space map [23, 25, 26].

Despite the importance of Imc to the signaling of the highest priority stimulus by the OTid [22, 23], its functional properties are not well understood [27, 28]. Recent work in barn owls has revealed the unusual multilobed structure of spatial RFs in the Imc, which has been shown to underlie its combinatorially optimized encoding of visual space [29]. In addition, Imc neurons have been shown to exhibit global inhibitory surrounds that may serve as a substrate for stimulus competition within the Imc [27, 30]. Here, we investigated in detail the properties of multisensory stimulus competition in, and the signaling of the most salient stimulus by, the Imc. Specifically, we examined how the responses of Imc neurons to two competing stimuli changed, as their relative strength (salience) was varied systematically. Our results demonstrate that the Imc is itself an active site of competition, as opposed to being either a passive conduit of inhibition to OT or simply reflecting activity in the OT. Imc neurons display signatures of stimulus competition that are quantitatively different from the OTid on average, and qualitatively distinct from those of individual, spatially aligned OTid sites recorded simultaneously.

## RESULTS

### Switch-like and gradual response profiles in the Imc to competing visual stimuli

To examine relative strength-dependent stimulus competition in the Imc, we recorded extracellularly the responses of Imc neurons in the barn owl, using a previously published competition protocol (Methods; [11, 31], Fig 1A). We presented a visual stimulus (S1) of fixed strength (loom speed; Materials and Methods) within the RF of a recorded neuron, and measured responses when a second, competing visual stimulus (S2) of varying strengths was presented far outside the RF (> 30 ° away; Fig. 1A; [11]. The responses obtained to the paired presentation of S1 and S2 using this protocol, collectively called competitor-strength dependent response profiles or CRPs (Fig. 1BD – bottom panels; [11], were compared with the responses to S1 presented alone (Fig. 1BD – top panels); the two types of trials were interleaved randomly.

**Figure 1.**
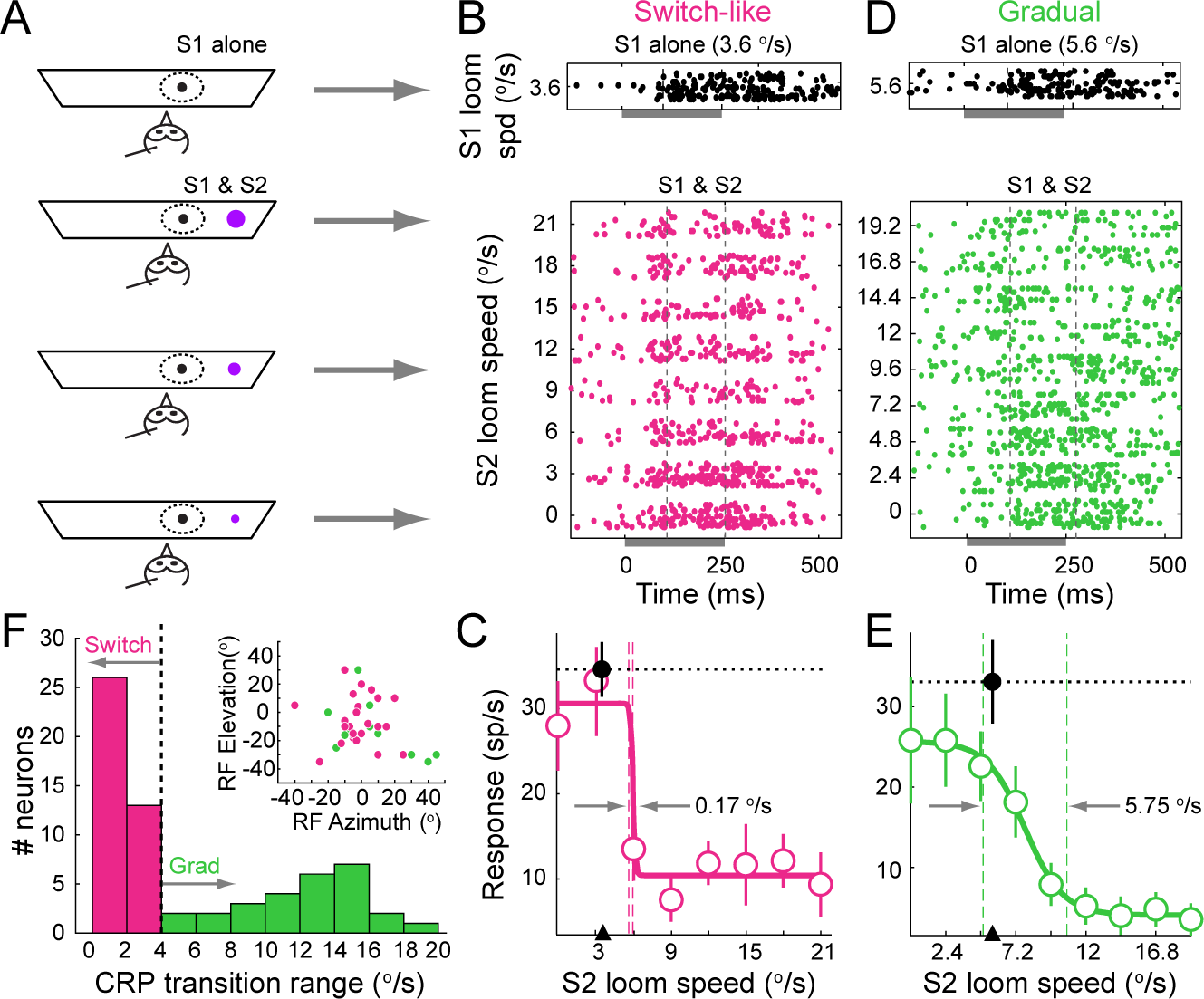
Switch-like and gradual response suppression of Imc neurons by a competing stimulus. **(A)** Schematic of experimental set up and stimulus protocol for measuring competitor strength-dependent response profiles (CRPs). Quadrilateral: tangent monitor; dashed oval: RF of recorded Imc neuron, black dot: S1 visual looming stimulus, magenta dots: S2 visual looming stimulus, sizes of dots denote loom speeds. **(B,C)** Responses of an example Imc neuron with switch-like CRP (data in magenta). (B) Rasters of spike responses to S1 alone (top; loom speed = 3.6 °/s), and to the paired stimuli (bottom) showing an abrupt increase in suppression with increasing S2 strength. Shaded box under x-axis represents stimulus duration (250 ms), dashed lines denote the time window (100-250 ms) during which response firing rates were calculated. (C) Response firing rates corresponding to rasters in B. Black data point (filled circle, and horizontal dotted line): response firing rate to S1 alone; mean ± s.e.m. Magenta data points (circles): response firing rates to paired presentation of S1 and S2 (i.e., CRP); mean ± s.e.m. Correlation coefficient of responses vs. S2 strength = −0.74 (p<0.05, Pearson correlation test). Solid line: best fitting sigmoid to the CRP, r^2^ = 0.95. Vertical dashed lines: transition range of this CRP (0.17 °/s; Materials and Methods). Black arrow head: Strength of S1 (3.6 °/s). **(D,E)** Responses of an example Imc neuron with gradual CRP (data in green). Conventions as in B,C. Loom speed of S1 = 5.6 °/s, correlation coefficient of responses vs. S2 strength = −0.94 (p<0.05, Pearson correlation test); r^2^ = 0.99 for the best fitting sigmoid; transition range = 5.75 °/s **(F)** Histogram of transition ranges of CRPs that exhibited a negative correlation with the strength of S2 (n=66 neurons/78 total). Vertical line: “cut-off” transition range of 4°/s (see Materials and Methods). The median strength of S1 was 7°/s with 95% CI of [6.3°/s, 7.7 °/s]; median distance of S2 from S1 = 43° with 95% CI of [40°, 46°]. Inset: RF locations (in double pole coordinates) of Imc neurons at which CRPs were recorded; colors correspond to whether the neurons had switch-like (magenta) or gradual (green) CRPs.

The majority of the recorded Imc neurons (66/78) exhibited CRPs that were negatively correlated with the strength of S2 (Fig. 1B-E; p<0.05, Pearson correlation test; Materials and methods). Of the remaining, one fraction showed CRPs with fixed response suppression, independent of the strength of S2 (1/78; Materials and Methods) and the rest were not affected by S2 (11/78; Materials and Methods).

Further examination of the negatively correlated CRPs revealed two distinct patterns of suppression based on how abruptly the responses transitioned from the maximum to the minimum value. In one set of CRPs, the majority of response suppression was expressed over a narrow range of S2 strengths, and in the other, the response suppression increased in graded, systematic way as a function of S2 strength. To quantify the abruptness of the response transition, we defined the CRP transition range as the range of S2 strengths over which the responses dropped from 90% to 10% of the maximum response (Materials and Methods). Following previously published convention [11]; Materials and Methods), CRPs with transition ranges narrower than 1/5^th^ the nominal range of S2 strengths (4°/sec), were referred to as being ‘switch-like’ (Fig 1B-C), while those with transition ranges broader than 4°/sec were referred to as being ‘gradual’ (Fig 1D-E).

Across the population of 66 Imc neurons that exhibited negatively correlated CRPs, the majority (39/66; 59%) of neurons exhibited switch-like responses and the rest (27/66; 40%) exhibited gradual CRPs (Fig. 1F). Switch-like and gradual CRPs were recorded at neurons both encoding for locations within the frontal portion of visual space (within ± 15° in azimuth and elevation), as well as in the more peripheral portion (Fig. 1F; inset), indicating that switch-like or gradual modulation of response suppression did not depend on the spatial location of the RF.

### Time course of stimulus competition in the Imc

The time course of response suppression was different between Imc neurons with gradual versus switch-like CRPs. For each neuron with a gradual CRP, we first calculated the instantaneous firing rate responses to S1 alone, and to the paired presentation of S1 and S2 for every strength of S2 (Materials and Methods). We then binned the relative competitor strength (S2-S1) values into five bins (Fig. 2A-H, columns), and for each bin, grouped across neurons the normalized instantaneous firing rate responses to S1 alone, and separately, to paired S1 and S2 (Fig. 2A vs. 2B, respectively; Materials and Methods). We quantified the emergence of response suppression for each relative strength bin by comparing the pooled responses to S1 and the pooled responses to paired S1 and S2 (Fig. 2C; gray vs. green, respectively) using a millisecond-by-millisecond running ANOVA procedure (Fig. 2D; Materials and Methods). The time-to-suppression (TTS) was defined as the first instant at which the responses diverged significantly (Fig. 2D; dashed vertical arrows; Materials and Methods).

**Figure 2.**
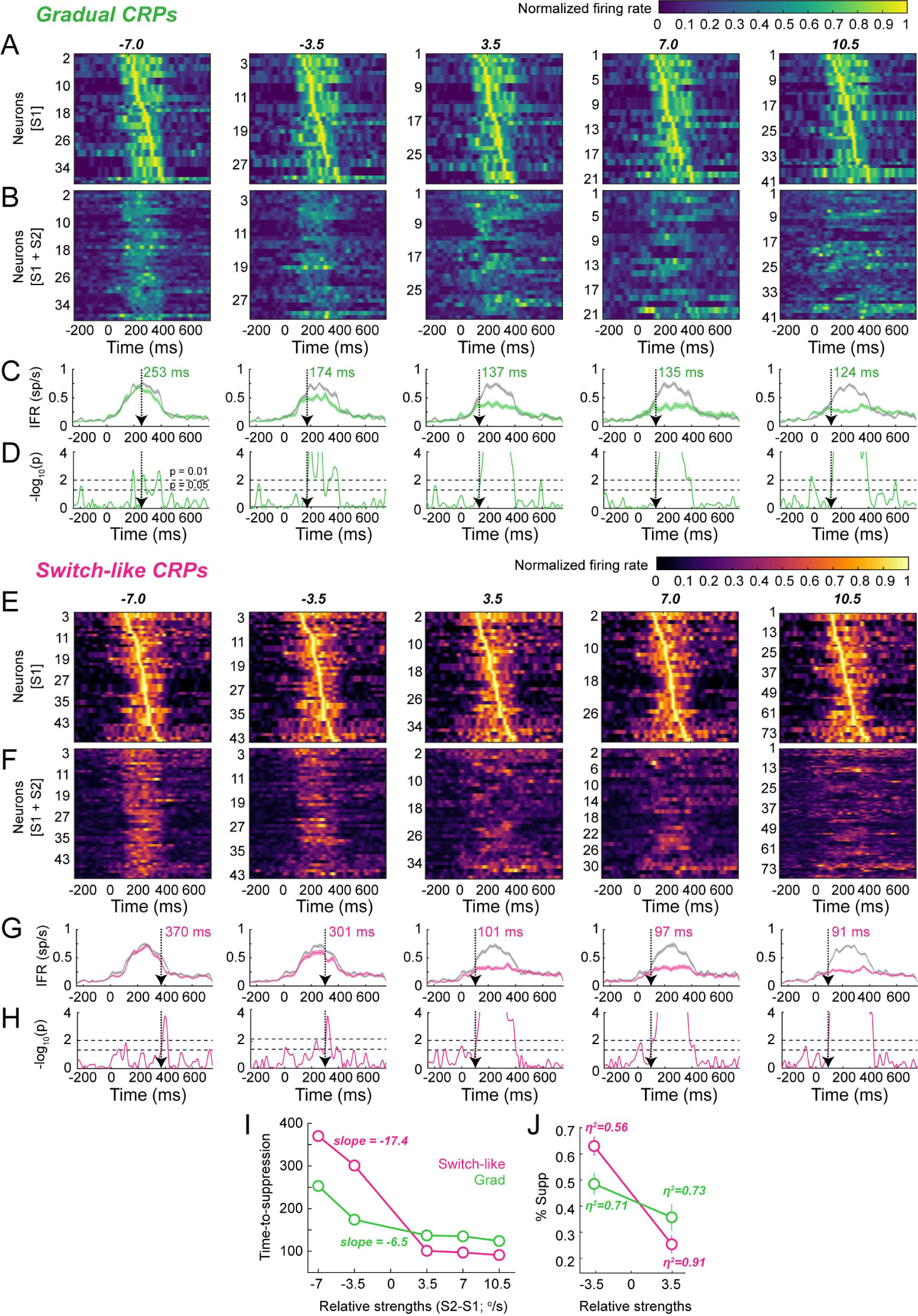
Time course of response suppression for Imc neurons with switch-like vs. gradual CRPs. **(A-D)** Analysis of response time courses of Imc neurons with gradual CRPs; columns - responses binned by the relative strength of betwen S1 and S2 (S2-S1). (A,B) Instantanous firing rates (IFRs) of neurons to S1 alone (A) or to S1 and S2 presented together (B), computed by smoothing PSTHs (1 ms time bins) with a Gaussian kernel (sd = 12 ms; Materials and Methods). For each neuron, IFRs are normalized by the peak firing rate of that neuron to S1 alone. Neurons in (A) are sorted by the half-peak firing rate; neurons in B are in the same order as in A. (C) Pooled average firing rates to S1 alone (gray) or to S1 and S2 (green) of all neurons within each bin. Translucent bands indicate s.e.m. (D) Time course of p-values obtained by performing ANOVA between the responses to S1 alone vs. to S1 and S2 at each millisecond. Vertical dashed arrows (and colored text): time-to-suppression (TTS); defined as the first instant at which responses to paired S1 and S2 diverge significantly from responses to S1 alone (Materials and Methods). Horizontal dashed lines: p-value thresholds used in determining TTS; lower line at p=0.05, upper line at p=0.01. **(E-H)** Same as A-D, but for Imc neurons with switch-like CRPs. **(I)** Comparison of TTS for neurons with switch-like (magenta) vs. gradual CRPs (green). **(J)** Plot of average amount of response suppression (± s.e.m) for switch-like vs. gradual neurons for the two relative strength bins on either side of the selection boundary. Text reports the effect size (eta^2^).

We found that among the neurons with gradual CRPs, there was a systematic reduction in the time-to-suppression as a function of relative strength: from 253 ms at relative strength = −7 °/s, to 124 ms at relative strength +10 °/s (Fig. 2I; green data; slope = −6.5, p < 0.05, linear regression). We repeated the above analysis for neurons with switch-like CRPs and found that the times-to suppression also decreased as a function of relative strength (from 370 ms to 91 ms), but with a much steeper slope than for neurons with gradual CRP (Fig. 2I; magenta data; slope = −17.4, p < 0.05, linear regression). There was, however, no systematic difference in the response latency of neurons with gradual versus switch-like CRPs (latency of response to S1 alone ̶ gradual: median = 136 ms with 95% CI [113,159]; switch-like: median = 110 ms with 95% CI [95 ms, 125 ms]; p > 0.05, sign test).

Notably, the times-to-suppression for switch-like CRPs were much longer than for gradual CRPs when S2 was weaker than S1 (Fig. 2I; magenta vs. green data at negative relative strength values), but flipped over to being shorter when S2 was stronger than S1 (Fig. 2I; magenta vs. green data at positive relative strength values). This resulted in a substantially large change in the time-to-suppression across the relative strength of zero (i.e., the ‘selection boundary’) for switch-like CRPs compared to gradual CRPs (switch-like: 200 ms drop from 301 ms to 101 ms vs. gradual: 37 ms drop from 174 ms to 137 ms).

We wondered if this flip in TTS values for switch-like vs. gradual CRPs across the selection boundary might be explained by the intrinsic difference in the shapes of switch-like versus gradual CRPs (Fig. 1C vs. 1E). Specifically, we asked if the systematic reduction in firing rates for gradual CRPs to paired S1 and S2 as a function of S2 strength (Fig. 1E) resulted in greater suppression of responses (than for switch-like CRPs) when S2 was just weaker than S1, but weaker suppression (than for switch-like CRPs) when S2 was just stronger than S1. The pooled population averages of instantaneous firing rates (Figs. 2C vs. G) suggested that this may be true. To test this explicitly, we quantified the amount of response suppression to paired S1 and S2 at relative strengths of −3.5 and +3.5 ̶ the two bins just on either side of the selection boundary. Indeed, we found greater suppression for gradual CRPs than switch-like CRPs at relative strength = −3.5, but weaker suppression at relative strength = +3.5 (Fig. 2J), consistent with the faster time-to-suppression for gradual CRPs at relative strength = −3.5 but slower time-to-suppression for gradual CRPs at relative strength = +3.5.

### Multisensory stimulus competition in the Imc

The occurrence of gradual and switch-like CRPs was not restricted to just the visual sensory modality. We measured “auditory” CRPs using a visual S1 (of fixed strength) and an auditory S2 (of varying strengths; S2_aud_). S2_aud_ stimuli were broadband noise bursts, and S2_aud_ ‘strength’ was varied by changing the binaural sound level (Materials and Methods).

We recorded auditory CRPs at 35 Imc neurons, and of these, 20 exhibited CRPs that were negatively correlated with the strength of S2_aud_ (Methods; p<0.05, Pearson correlation test). Further examination of the responses of these neurons revealed two distinct patterns of suppression as a function of strength of S2_aud_: gradual or switch-like (Fig 3B-E). Switch-like auditory CRPs were defined as those for which the transition range was narrower than 9 dB (1/5^th^ the full range of S2_aud_ strengths; just as in the case of visual CRPs, and consistent with previously published literature – [11, 32]. Of the 20 neurons with correlated CRPs, 10 were found to be gradual and the rest, switch-like (Fig 3D). Thus, consistent with Imc’s role in enabling within as well as cross-sensory stimulus competition in the OTid [22], Imc neurons themselves exhibit signatures of multisensory stimulus competition [30].

**Figure 3.**
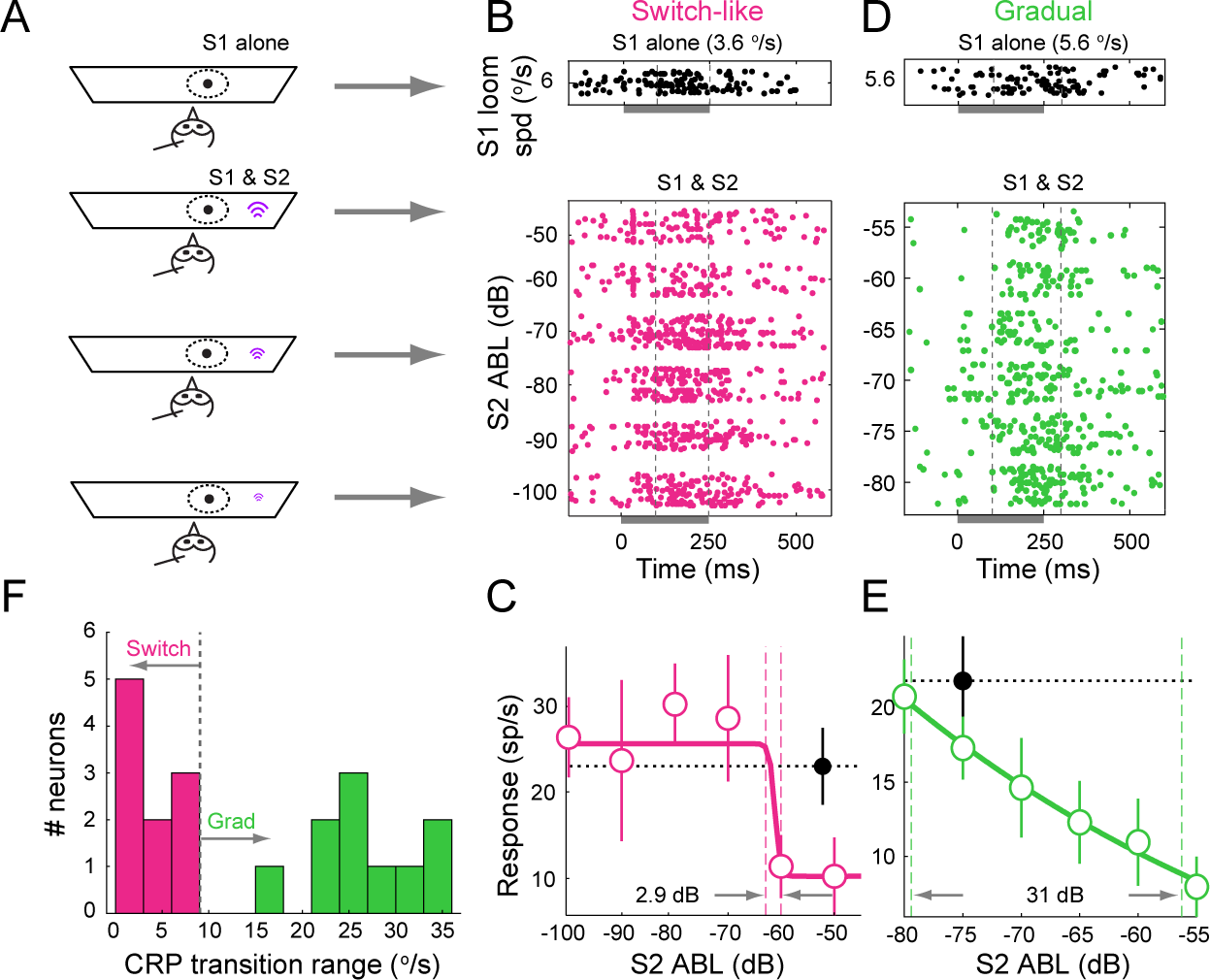
Switch-like and gradual response suppression of Imc neurons by an auditory competitor. **(A)** Schematic of experimental set up and stimulus protocol for measuring auditory CRPs (conventions as in Fig. 1A). Black dot: S1 visual looming stimulus. Magenta sound symbol: S2 auditory stimulus (S2_aud_); size of symbol represents binaural intensity of S2_aud_. **(B-C)** Switch-like auditory CRP measured at an example Imc neuron. (B) Response rasters. Dashed lines: time window (100-250 ms) during which response firing rates were calculated. S1 loom speed = 6 °/s. (C) spike counts. Correlation coefficient of responses vs. S2_aud_ strength = −0.76 (p<0.05, Pearson correlation test); r^2^ = 0.94 for the best fitting sigmoid, transition range = 2.9 dB. All other conventions as in Fig. 1BC. **(D-E)** Gradual auditory CRP measured at an example Imc neuron. (D) rasters. Dashed lines: time window (100-300 ms) during which response firing rates were calculated. S1 loom speed = 5.6 °/s. (E) spike counts. Correlation coefficient of responses vs. S2_aud_ strength = 0.99 (p < 0.01, Pearson correlation test), r^2^ = 0.99 for the best fitting sigmoid, transition range = 31 dB. All other conventions as in Fig. 1DE. **(F)** Histogram of transition ranges of CRPs recorded in Imc that exhibited a negative correlation with the strength of S2_aud_ (n=20 neurons/35 total). Vertical line: “cut-off” transition range of 9 dB (Materials and Methods). The median strength of S1 was 5.6°/s with 95% CI of [5.1°/s, 6.1 °/s].

### Imc signals the strongest stimulus accurately and flexibly

The observation of abrupt response suppression in switch-like CRPs led us to ask if strength of S2 at which the transition occurred from high to low response values was meaningful. Because this transition was also well-defined (albeit less so) in gradual CRPs, we asked this question more generally of switch-like as well as grdual CRPs in the Imc. Specifically, we wondered if the strength of S2 at which the transition occurred was related to the (fixed) strength of the stimulus inside the RF, S1. (Because this comparison is only meaningful when both stimuli are of the same sensory modality, we restricted our analysis to visual CRPs.) To this end, we first defined as the CRP ‘transition value’ the strength of S2 at which the responses were half-way between the maximum and minimum values, and quantified it as the midpoint of the transition range (Fig. 4A; Materials and Methods). We then compared the transition value of each CRP to the strength of S1 used to measure the CRP, and defined this difference as the CRP ‘bias’ (bias = transition value – strength of S1; Fig. 4A).

**Figure 4.**
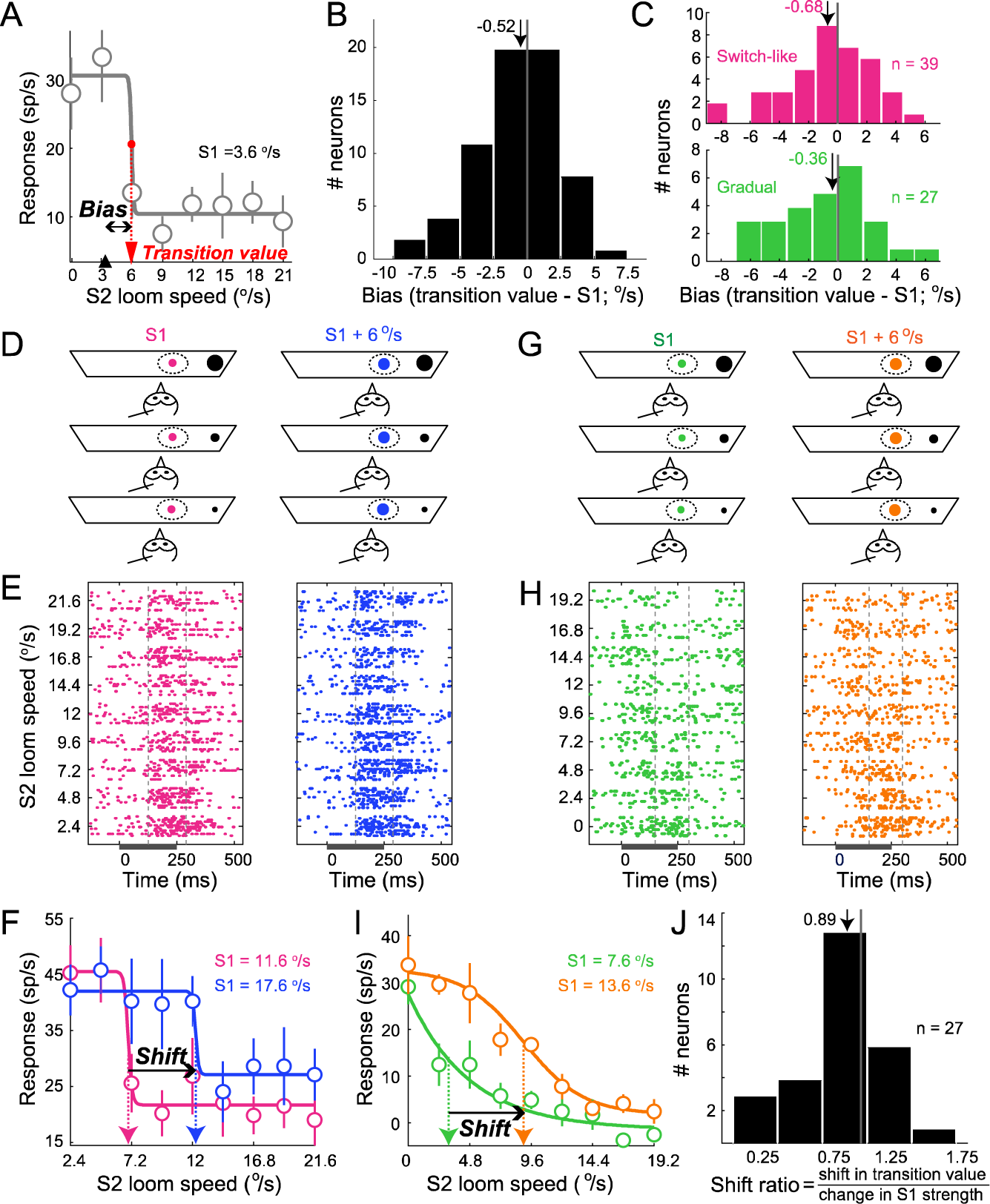
Dependence of CRP transition value on S1 strength for Imc neurons. **(A)** Definition of CRP transition value and CRP bias for switch-like visual CRP in Fig. 1C (see Materials and Methods). S1 = 3.6 °/s; transition value = 5.9 °/s; bias = 2.3°/s. **(B)** Distribution of CRP bias for visual CRPs; n= 66 Imc neurons. Median = −0.52, p = 0.39; sign test vs. 0. **(C)** Distributions of CRP bias, separately for Imc neurons with switch-like CRPs (top panel) and gradual CRPs (bottom panel). Switch-like: median bias = −0.68, p = 0.52, sign test against 0 followed by Holm-Bonferroni correction for multiple comparisons, n=39 neurons; gradual: median bias = −0.36, p = 0.7, sign test against 0 followed by Holm-Bonferroni correction for multiple comparisons, n=27 neurons. **(D)** Schematic of experimental protocol to measure two CRPs with S1 stimuli of two different strengths, indicated by magenta and blue dots (D). S1 =11.6 °/s; magenta data, and S1 = 17.6 °/s; blue data. **(E, F)** Two CRPs (shown in magenta and blue, respectively) measured using protocol in D at an example neuron with switch-like CRP. E – rasters, F – firing rate plots, conventions as in Fig. 1. Transition values: 6.95 °/s (when S1 = 11.6; magenta data) and 12.3°/s (when S1 = 17.6 °/s; blue data). Shift ratio (shift in transition value / change in S1 strength) for this example neuron = 0.88. **(G, H, I)** Schematic (G), and two CRPs (H,I) measured at an example neuron with gradual CRP. Transition values: 3.22 °/s (when S1 = 7.6 °/s; green data) and 9.01 °/s (when S1 = 13.6 °/s; orange data). Shift ratio (shift in transition value / change in S1 strength) for this example neuron = 0.96. **(J)** Distribution of the shift ratio across n= 27 Imc neurons. Median shift ratio = 0.89, p = 0.26, sign test against 1.

Across the population of Imc neurons with correlated visual CRPs (n=66), we found neurons with a range of CRP biases (Fig. 4B). Some had a negative bias, indicating that these CRPs transitioned from a high to a low value when S2 was less than S1, and others had a positive bias, indicating that those CRPs transitioned from a high to a low value when S2 was greater than S1. However, across the population, the CRP bias was distributed around zero (Fig. 4B; median CRP bias = −0.52, p= 0.39, sign test against 0). This indicated that on average, Imc neurons responded at a high level when S1 (RF stimulus) was the strongest stimulus, but transitioned to responding at a low level when S1 was no longer the strongest stimulus, i.e., just when S2 exceeded S1 in strength. This was true separately both for switch-like as well as for gradual CRPs (Fig. 4C; top and bottom panels, respectively; switch-like: median bias = −0.68, p = 0.52, sign test against 0, n=39 neurons; gradual: median bias = −0.36, p = 0.7, sign test against 0, n=27 neurons).

These results indicated that Imc may signal the strongest of the competing stimuli without any bias, i.e., ‘accurately’, and suggested the interesting possibility that transition values of Imc CRPs are not fixed quantities, but are coupled ‘flexibly’ to the strength of S1. To test this hypothesis that CRP transition values depend on the strength of S1, we measured two CRPs for each of a subset of Imc neurons. One CRP was measured with a weaker S1, another with a stronger S1 (S1 + 6 °/sec), with S2 varying over the same range of loom speeds in both cases (Fig. 4DG; [11]). The stimuli corresponding to the two CRPs were presented in a randomly interleaved manner. We found that the CRP transition value shifted with S1 strength in the predicted way: a stronger S1 produced a right shifted CRP (Fig. 4EF – data from example neuron with switch-like CRP showing a right shift in transition value; Fig. 4HI – data from example neuron with gradual CRP showing a right shift). Across the population of tested neurons, we found that the magnitude of the shift in CRP transition value matched on average, the magnitude of the change in S1 strength (Fig. 4H; median shift ratio (shift in transition values / change in S1 strength) = 0.89, p = 0.26, sign test against 1, n=27 neurons).

Taken together, these results established that Imc neurons report dynamically, an online comparison between the strengths of the two competing stimuli. They signal accurately (with average bias indistinguishable from 0), the strongest of the two competing stimuli, and do so flexibly (with transition values coupled to the strength of S1).

### Comparison of signatures of stimulus competition in the Imc and OTid

The findings of switch-like and gradual CRPs in the Imc, as well as accurate and flexible signaling of the strongest stimulus, parallel previous findings in two other key nuclei in the midbrain selection network, namely the OTid [11–13] and the cholinergic Ipc [32]. These nuclei are interconnected: Imc receives focal input from layer 10 of the OT (OT_10_), but projects back broadly across the OTid as well as the Ipc [4], and Ipc receives focal input from OT_10_ and projects back focally to the OT [24], serving to amplify OTid responses [23, 25]. Additionally, Imc is the source of competitive inhibition in the OTid and Ipc: focal inactivation of Imc abolishes competitive interactions in both areas [22, 23]. Considering this interconnectedness, we were interested in whether signatures of stimulus competition in the Imc simply reflect computations occurring elsewhere (for instance, in the OT), or if the Imc, itself, serves as an active site at which computations related to stimulus competition occur. We focused on the comparison between Imc and OTid (rather than Ipc as well) because the OTid is the “output” hub of the midbrain selection network, involved in controlling behavior as well as relaying information from the midbrain to cortical areas [33, 34].

To address this question, we compared metrics of stimulus competition in Imc and OTid. In order to minimize the impact of any idiosyncratic differences in spike sorting, selection of count windows, or other analysis choices on this comparison, we measured CRPs in the OTid as well (Materials and Methods; [11]).

First, we compared the accuracy with which Imc vs. OTid neurons, on average, signal the strongest stimulus. We found that the CRP bias distributions were not distinguishable (Fig. 5A; red vs. blue data; p > 0.05, sign test), indicating that Imc and OTid neurons signaled the strongest stimulus with comparable accuracy.

**Fig. 5:**
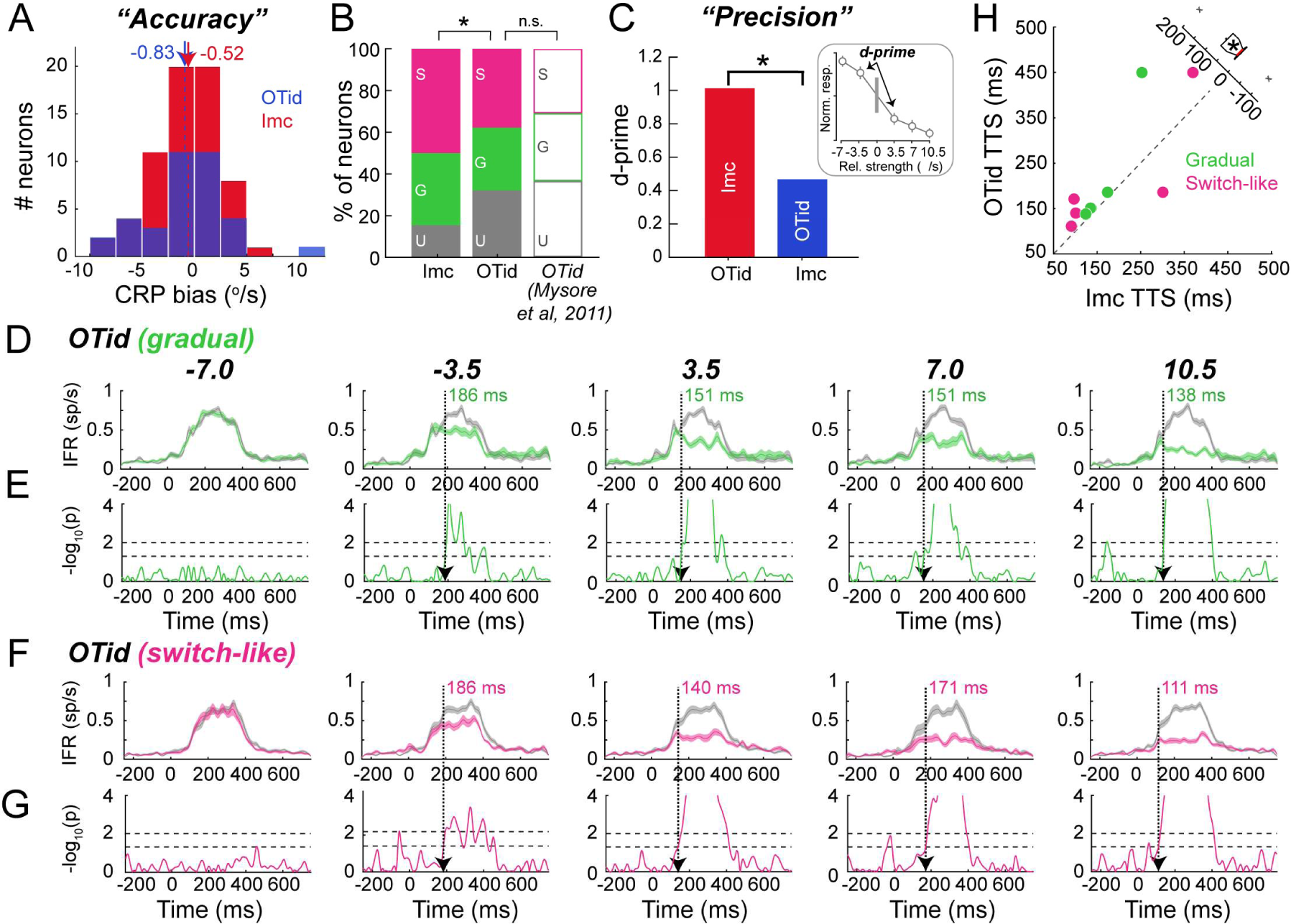
Imc signals stimulus competition with greater precision, and earlier than OTid. **(A)** Distributions of CRP bias for OTid neurons (blue; n=36 neurons with correlated CRPs; median=−0.83; p>0.05; sign test against 0 and Imc neurons (red; data reproduced from Fig 4B; n=66; median = −0.52). No significant difference between Imc and OTid medians (p>0.05, sign test after Holm-Bonferroni correction for multiple comparisons). **(B)** Proportions of switch-like (S), gradual (G), and uncorrelated (U) CRPs measured in Imc (stacked bar on left; n=78 neurons at which CRPs where measured) and OTid (middle bar; n= 53 neurons at which CRPs were measured). Previously published OTid proportions (n=169 neurons at which CRPs were measured; data adapted from [11]) are also shown for completeness (right bar; unfilled). ‘*’ (‘n.s.’): p<0.05 (p>0.05), chi-squared test between Imc and OTid proportions measured here followed by correction for multiple comparisons. **(C)** Comparison of d-prime (Materials and Methods) computed from pooled CRP responses measured in OTid (blue) vs. Imc (red). ‘*’: p<0.05, permutation test (Materials and Methods). Inset: Schematic of pooled CRP responses across neurons illustrating d-prime computation across the selection boundary (vertical gray line). **(E)** Pooled instantaneous firing rates of OTid neurons with gradual CRPs in response to S1 alone (grey) or paired S1 and S2 (green), binned into five relative strength bins (columns). **(F)** Millisecond-by-millisecond running ANOVA to determine time-to-suppression (vertical dashed line): the first instant at which responses to paired S1 and S2 diverge significantly from responses to S1 alone (Materials and Methods). OTid responses never diverge significantly for the first relative strength bin (S2-S1 = −7 °/s; left-most panel). **(G,H)** Same as E,F, but for OTid neurons with switch-like CRPs (magenta data); OTid responses never diverge significantly for the first relative strength bin (S2-S1 = −7 °/s; left-most column). **(I)** <u>Left:</u> Scatter plot of TTS measured in Imc versus in OTid. Dots: (Imc, OTid) TTS pairs for the different relative strength bins; magenta data – switch-like CRPs; green data – gradual CRPs. For plotting purposes, TTS values corresponding to cases in which the responses to paired S1 and S2 never diverged from those to S1 alone, are coded as 450 ms. Dashed line: line of equality. <u>Right:</u> Boxplot of differences between OTid and Imc TTS values; p<0.05, sign test against 0.

Next, we compared the relative proportions of neurons that exhibited switch-like, gradual or uncorrelated CRPs in the Imc vs. OTid (Fig. 5B). The proportions were different between Imc and OTid: (Imc: n=78 neurons; 50% switch-like, 34.6% gradual, and 15.4% uncorrelated CRPs; OTid: n=53 neurons; 37.7% switch-like, 30.2% gradual, and 32.1% uncorrelated CRPs; p<0.01, chi-squared test followed by Holm-Bonferroni correction for multiple comparisons). Indeed, the proportions of CRP types measured in the OTid in this study were not significantly different from those previously published, confirming the veracity of our results (OTid published data, Mysore et al 2011, n=169 neurons, 30.2% switch-like, 33.1% gradual, and 36.7% uncorrelated CRPs; p=0.72, chi-squared test between measured and published OTid followed by Holm-Bonferroni correction for multiple comparisons).

The larger fraction of switch-like CRPs in the Imc than the OTid (50% Imc vs. 37.7% OTid) suggested that Imc ensembles may be able to signal the strongest stimulus more categorically than the OTid [12], or in other words, that the signaling of the strongest stimulus may be more “precise” in the Imc (Fig. 5C). To test this directly, we quantified a metric of categorical signaling: the discriminability (d’) across the selection boundary (relative strength =0 °/s; S1 = S2[11]; Materials and Methods). To compute this metric, we pooled CRP responses across all recorded Imc neurons (gradual, switch-like and uncorrelated CRPs) binned into 5 relative strength bins, and then calculated the d-prime between the pooled responses in the relative strength bin of −3.5 °/s versus the relative strength bin of +3.5 °/s (straddling the boundary; Fig. 5C-inset; Materials and Methods). We repeated this across all OTid neurons. This approach allowed us to estimate the ability of a downstream neuron of Imc (or OTid; ideal observer) to decode the strongest stimulus from population activity in the Imc (OTid). We found that d’ across the selection boundary (S1=S2) was nearly twice as high in the Imc as the OTid (Fig. 5C; Imc =1.02, OTid = 0.46; p < 0.01, permutation test). This result established that the Imc signaled the strongest stimulus more categorically (with greater precision) than the OTid.

Finally, we compared the time course of stimulus competition in the Imc vs. OTid. To this end, we computed the instantaneous firing rates of OTid neurons to S1, and to paired S1 and S2, for each CRP and each strength of S2 (Materials and Methods). Following the procedure employed for analyzing Imc time courses, we binned paired S1+S2 responses into 5 bins. Within each bin, we pooled the instantaneous firing rates across switch-like (and separately, across gradual) OTid neurons (Fig. 5DF; Materials and Methods), and compared the pooled responses to S1 alone with those of S1+S2 using a millisecond-by-millisecond ANOVA procedure. We quantified separately for OTid neurons with gradual CRPs (Fig. 5E) and switch-like CRPs (Fig. 5G), the time-to-suppression at the different relative strength bins (Fig. 5EG; Materials and Methods). We found that times-to suppression were consistently faster in the Imc than the OTid (Fig. 5H; Imc faster than OT by median value of 18 ms; p<0.05, sign test).

Taken together, these quantitative differences in precision and time course of responses to paired S1 and S2 indicate that the Imc is itself a site where computations related to stimulus competition occur, and also that the presence of a competitor is signaled first in the Imc followed by the OTid.

### Simultaneous, paired measurements of stimulus competition in the Imc and OTid

As a final step, we wanted to go beyond comparisons between Imc and OTid populations recorded independently, and compare directly signatures of competition in simultaneously recorded Imc and OT sites encoding for the same portion of sensory space. This approach can reveal how stimulus competition in the two brain areas unfolds at the same time during exposure to the same competing stimuli.

To this end, we first positioned an electrode in the Imc, mapped the RF, and then positioned a second electrode in the OTid such that the OTid RF overlapped with the Imc RF (Fig. 6A; spatially ‘aligned’ OTid and Imc RFs - dashed ovals). We simultaneously recorded OTid and Imc responses while presenting S1 and S2 per the CRP stimulus protocol: S1 was presented within the overlapping portion of the RFs, and S2 was presented far outside (> 30° away; Fig. 6A – S1 and S2; both were visual looming dots; Materials and Methods).

**Figure 6.**
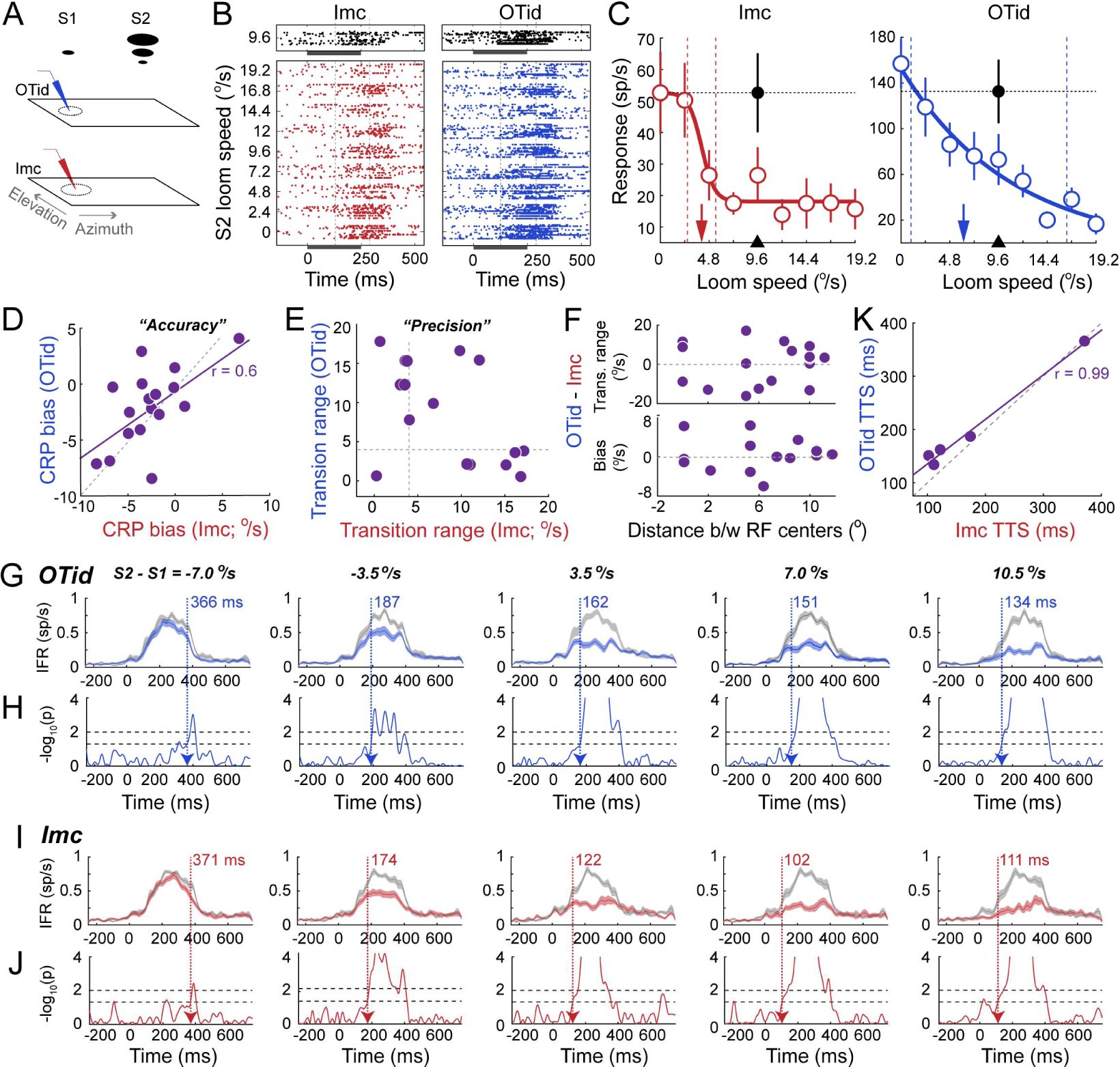
Signatures of stimulus competition at simultaneously recorded, aligned Imc and OTid sites. **(A)** Schematic of experimental set-up for simultaneous, paired recordings in Imc and OTid. S1 and S2: stimulus protocol for measuring CRPs; colored icons to the left: recording electrodes, positioned in OTid (blue) and Imc (red); dashed ovals: RFs of recorded site. **(B)** Raster plots of responses for example paired, aligned Imc (left panels) and OTid (right panels) sites; distance between RF centers = 8° (Materials and Methods). Top panels: responses to S1 alone; bottom panels: responses to S1 and S2 presented together. Strength of S1 = 9.6 °/s. All other conventions as in Fig. 1B. **(C)** Firing rates responses corresponding to rasters in B. Conventions as in Fig. 1C. CRP correlation values: Imc = −0.88, p<0.05; OTid = −0.99, p<0.05. CRP transition ranges: Imc = 2.8 °/s (switch-like CRP); OTid = 15.3 °/s (gradual CRP). CRP transition values: Imc = 4.1 °/s; OTid = 6.2 °/s. **(D)** Scatter plot of CRP biases measured at paired Imc and OTid sites (n = 17 pairs; average distance between RF centers = 6.3 ° +/−0.95 °). Dashed grey line: line of equality. Solid magenta line: best fit line to data; Pearson’s ρ = 0.6, p=0.01. **(E)** Scatter plot of CRP transition ranges measured at paired Imc and OTid sites. Dashed gray lines: mark cut-off value of transition ranges (4 °/s) for switch-like vs. gradual CRPs. **(F)** Plot of difference in CRP transition ranges (OTid – Imc; top panel) or CRP biases (OTid – Imc; bottom panel) for paired Imc-OTid sites as a function of distance between Imc and OTid RFs. Pearson’s ρ = 0.03, p=0.9 (CRP transition ranges); ρ = 0.07, p=0.8 (CRP biases). **(G-J)** Response time courses for OTid (G; blue) and Imc sites (I; red) recorded simultaneously; shown are the pooled averages across OTid sites, and separately, across Imc sites (conventions as in Fig. 2). Results from the millisecond-by-millisecond AVOVA are also shown for OTid (H; blue) and Imc sites (J; red), respectively. Columns: Relative strength bins. IFR: Instantaneous firing rate. Horizontal dashed lines: p-value cut offs (0.05, lower line, and 0.01, upper line). Vertical dashed arrows (and colored text): time-to-suppression (TTS; Materials and Methods). **(K)** Scatter plot of TTS measured at aligned Imc vs. OTid sites recorded simultaneously. Each dot: TTS pair for a different relative strength bin; data correspond to the colored numbers in G,J. Dashed line: line of equality. Pearson’s ρ = 0.99, p<0.05.

Responses from an example pair of simultaneously recorded, aligned Imc and OTid sites (distance between RF centers = 8°; Materials and Methods) showed that the nature of the CRP was different in the two areas (Fig. 6BC: switch-like in Imc and gradual in OTid). However, both CRPs exhibited negative bias (Fig. 6C: vertical arrows to the left of black arrowheads).

We quantified these properties for each aligned site-pair in our population (n=26 pairs) for which *both* Imc and OTid CRPs were negatively correlated with the strength of S2 (n=17 pairs). Across these 17 pairs of Imc-OTid sites (average difference in centers of RF = 6.3 ° +/−0.95°), we found that CRP biases in Imc were not different from those in OTid (Fig. 6D; p>0.05, sign test of TTS differences between Imc and OTid). Notably, CRP biases at paired Imc and OTid sites were positively correlated (Fig. 6D; Pearson’s ρ = 0.6, p=0.01). By contrast, CRP transition ranges at paired Imc and OTid sites were uncorrelated (Fig. 6E; Pearson’s ρ = −0.37, p = 0.15). These results regarding CRP biases and transition ranges did not depend on the degree of alignment between the paired Imc and OTid sites (Fig. 6F; bias vs. alignment, Pearson’s ρ = 0.07, p=0.8; transition ranges vs. alignment, Pearson’s ρ = 0.03, p=0.9). Thus, for Imc and OTid sites encoding for the same portion of sensory space, accuracy of signaling the strongest stimulus was correlated, but precision of the signaling was not.

Finally, we examined the speed at which paired Imc and OTid sites signaled the presence of a competing stimulus. We compared the time course of response suppression by calculating (as before) the time-to-suppression within each relative strength bin for OTid sites (Fig. 6GH) as well as paired Imc sites (Fig. 6IJ). The times-to-suppression at paired Imc and OTid sites were highly correlated (Pearson’s ρ =0.99, p<0.05, correlation test), with Imc sites signaling the presence of the competitor earlier than paired OTid, consistent with our findings from independent recordings (Fig. 6K; best fit line has positive intercept, intercept = 52.8, 95% CI [16.2,89.4], with slope not different from 1, slope = 0.84, 95% CI [0.66,1.01], indicating that OTid TTS are significantly above line of equality).

## DISCUSSION

This study elucidates the properties of multisensory, salience - dependent stimulus competition in a pivotal nucleus in the midbrain selection network in vertebrates, namely the Imc [4, 5, 7, 8, 35]. This small group of GABAergic neurons [4, 29], which supplies inhibition in a combinatorial manner to all parts of the OT space map [4, 29], serves a critical function: without it, competitive interactions and selection in the OTid are abolished [22, 23]. Considering the critical role of the intermediate and deep layers of the SCid in target selection for spatial attention [9, 10], the Imc appears to occupy a spot of central importance within the vertebrate midbrain selection network.

One manner in which the Imc might control competition and selection in the OTid is by serving as a passive relay of inhibition, simply flipping the sign on the input excitatory drive from OT_10_. Together with Imc’s anatomical projection patterns, this implementation would allow Imc to facilitate computations in the OTid. However, another possibility is that the Imc is itself a site at which computations relating to stimulus competition occur actively, i.e., one at which information about competing stimuli is compared, with this processed information then being relayed to downstream targets (Ipc and OTid). Our results directly support the latter hypothesis.

We found that most Imc neurons (~85%) responded to a visual RF stimulus (S1) with decreasing firing rates as the strength of a distant visual competitor (S2) was systematically increased. The responses transitioned from a high to a low value in an abrupt (switch-like) manner in the majority of these cases (60%), and gradually in the others. Notably, the strength of S2 at which the transition from high-to-low responses occurred was coupled to the strength of S1, and was, on average equal to it. These results demonstrated that Imc neurons perform an online comparison of the strengths of the competing stimuli, and signal the strongest one. The large proportion (60%) of switch-like response profiles resulted in the Imc signaling categorically the strongest stimulus – we have shown in previous work that a population of neurons in which 30% or more exhibit switch-like competitive response profiles produces categorical signaling at the level of the entire ensemble [11].

Imc neurons also exhibited multisensory stimulus competition. When Imc was tested with competing stimuli of different sensory modalities, we found qualitatively similar results to when both stimuli were visual. These results indicated that Imc signals the strongest stimulus, independently of the sensory modalities. Notably, building off of findings that the average transition value of Imc response profiles is equal to the strength of S1, in the auditory case, the average transition value (−71 dB) across neurons presents an estimate of the binaural level of an auditory competitor that the Imc deems to be equivalent in strength to the average loom speed of S1 (6.9 °/s).

Our results also showed distinct time courses of responses for neurons with switch-like versus gradual CRPs. When S2 was weaker than S1, switch-like neurons signaled the presence of a competitor later than gradual neurons, but when S2 was stronger than S1, switch-like neurons were faster. This potentially puzzling ‘flip’ was accounted for by the intrinsic differences in the shapes of switch-like versus gradual neuron responses, which resulted in greater amounts of response suppression for gradual CRPs when the competitor was weaker than the stimulus in the RF (S2 < S1) but greater amount of suppression for switch-like CRPs when the competitor was stronger (S2 > S1), indicating that compared to neurons with gradual CRPs, neurons with switch-like CRPs quickly and effectively reflect response suppression when a competing stimulus is the stronger one. These differences are also potentially consistent with circuit mechanisms necessary for producing switch-like response profiles [36].

The signatures of competition in the Imc are quantitatively and qualitatively different from those in the OTid. Imc and OTid neurons both signaled the strongest stimulus accurately, with almost no bias in estimating when the two competing stimuli were equal in strength. However, with respect to another key aspect of stimulus competition, namely, the precision with which neurons signal the strongest stimulus ̶ either in a binary-like, explicitly categorical manner, or in a more analog, gradual manner ̶ Imc differed quantitatively from the OTid: it signaled the strongest stimulus much more accurately (2x better). These results first came to light from data collected independently (in different experiments) in the Imc and OTid, but using the same experimental set-up, stimulus protocols and analysis pipelines. Subsequently, paired simultaneous recordings in spatially aligned portions of the Imc and OTid not only confirmed these findings, but extended them. They revealed that some aspects of stimulus competition occurred in a coordinated manner in portions of Imc and OTid that encode for the same region of sensory space, i.e., that are active at the same time in a bird experiencing the competing stimuli. Specifically, the bias of neurons in estimating whether a competing stimulus was weaker or stronger than their RF stimulus was highly correlated. This suggests the presence of a shared or mutually dependent mechanism in setting the bias of competition. By contrast, the precision with which neurons signaled the strongest stimulus was not correlated between Imc and OTId neurons encoding for overlapping regions of sensory space. This suggests the presence of independent mechanisms in these two areas involved in setting the precision of competition, providing further support for Imc being an active, independent locus of competition, rather than a simple inhibitory relay.

Analysis of response time courses in both the separate as well as paired Imc-OTid experiments demonstrated that the Imc reports stimulus competition and signals the strongest stimulus earlier than the OTid. Considering that the Ipc, the other key nucleus in the midbrain selection network is also a downstream target of the Imc (just as the OTid is), it is plausible that the Ipc will also be slower than the Imc at signaling the strongest stimulus, just as the OTid is (but needs to be tested). Consequently, the Imc emerges as the site within the heavily interconnected midbrain selection network at which stimulus competition is potentially resolved first. In any case, the finding above, along with Imc’s categorical signaling and anatomical connectivity, reveals that the Imc sends differentially (categorically) enhanced competitive inhibition to OTid and Ipc sites encoding the weaker versus the stronger competing stimuli.

In summary, not does Imc actively perform computations of relative strength dependent stimulus competition, and does so earlier than the OTid, some aspects of these computations differ qualitatively from paired OTid sites. Overall, the Imc is more categorical in its signaling of the strongest stimulus.

The mechanism by which competition within the Imc may occur is yet to be demonstrated directly. Clues come from modeling and slice experiments, respectively predicting [36] and demonstrating [37] the presence of long-range inhibitory projections between Imc neurons. These projections could serve to deliver the competitive inhibition [38] necessary for the functional signatures reported here.

Considering that the midbrain selection network is conserved across vertebrate species [4–8], our findings have the power to guide understanding of the midbrain mechanisms underlying spatial selection in other vertebrates. More generally, these findings have implications for unpacking how key computations underlying various forms of selection such as perceptual categorization, value-based decision-making, etc., may be implemented in neural circuits [39]. The ubiquity of selection in adaptive behavior, coupled with our current lack of understanding of the precise neural mechanisms that underpin it, highlight the importance of the barn owl midbrain selection circuit as a gateway for generating hypotheses about viable circuit solutions for selection and decision-making in general.

## MATERIALS AND METHODS

### Neurophysiology

Eleven adult barn owls (Tyto alba; male and female; shared across different studies) were used for electrophysiological recordings. Birds were group housed in an aviary with a 12hr/12hr light/dark cycle. All protocols for animal care and use followed approval by the Johns Hopkins University Institutional Animal Care and Use Committee, and were in accordance with NIH guidelines for care and use of laboratory animals. All experimental and surgical procedures followed previously published methods [30].

Briefly, owls were anesthetized with isoflurane (2%) and a mixture of nitrous oxide and oxygen (45:55) on experiment days and head-fixed in a sound-attenuating booth. The head-fixation was calibrated following published procedures [40] such that the dorsolateral tip of the pecten oculi structures within the eyes were positioned at 7° above the horizon and approximately 25° lateral to the vertical midline. Isoflurane was ceased after birds were secured. We recorded from single and multiunit sites in the Imc and OT either epoxy-coated tungsten microelectrodes (A-M Systems, 5 MΩ at 1kHz). Recording sites in the intermediate and deep layers of OT (OTid) and the Imc were targeted on the basis of stereotaxic coordinates (from prior experiments) and verified on the basis of established neural signatures as described elsewhere [11, 16, 22, 29, 30]. For Imc recordings, an electrode was positioned to enter the brain at a medial-leading angle of 5°. At a subset of sites, a second electrode was lowered into OTid to make dual recordings simultaneously. Upon positioning the recording electrode(s), nitrous oxide was turned off for the duration of the data collection in some experiments; previous work has established no effect of nitrous oxide tranquilization on neural responses to competition protocols in the midbrain network [11]. Spike times were recorded using Tucker-Davis hardware and custom MATLAB software. Multiunit spike waveforms (OTid and Imc) were sorted into single neurons using the Chronux spike-sorting toolbox [41]. The quality of the sorted neurons were assessed visually, and additionally subjected to an F-test to determine whether or not each neuron was well-isolated from other neurons recorded within the same multiunit site [29]; only well-isolated neurons were retained.

### Stimuli

Visual stimuli were presented as black, fixed contrast, expanding looming dots on a grey background on a 65” monitor. Looming dot stimuli were used as these reliably evoke strong responses in OT and Imc [11, 16]. The strength of a looming stimulus was defined by its loom speed, with faster loom speeds typically evoking greater responses; the typical range of loom speeds used was 0°/s to 20°/s. Locations of visual stimuli were defined by double pole coordinates relative to the midsagittal plane for azimuth or the visual plane for elevation [40]. Auditory stimuli were presented as broadband noise bursts with equalized amplitudes delivered binaurally through earphones. Sounds were filtered with a head-related transfer functions (HRTF) of a standard barn owl [16]. Strengths of auditory stimuli were defined by the auditory binaural level (ABL). Visual stimuli were generated using custom MATLAB scripts and psychtoolbox (PTB-3; [42, 43]), and auditory stimuli were generated using custom MATLAB scripts and Tucker Davis Technologies hardware.

Two-dimensional receptive fields (RFs) were collected by presenting a stimulus at various azimuthal and elevational locations. These stimuli were either a single looming dot of fixed strength or a stationary dot (radius 3°) moving at a 45° angle over 3°. For RF measurements, stimuli were presented for 5-7 repetitions, with a duration of 250 ms each and an inter stimulus interval of 1000-1500 ms. Spatial locations at which a single stimulus elicited higher firing rates compared to baseline were deemed to constitute neuron’s spatial RF, and were used to estimate RF extent (half-max-width) and center (weighted average of RF locations).

Stimulus competition protocols involved the presentation of a visual stimulus (S1) of fixed strength inside the RF, either by itself, or with a second stimulus (either visual or auditory; S2_vis_ or S2_aud_, respectively) of varying strengths presented at a distant location (typically 30° away from S1). The resulting responses from paired S1 and S2 presentation were collectively called competitor-strength dependent response profiles or CRPs [11]. Stimuli were presented for 10-15 repetitions, with a duration of 250 ms each and an inter stimulus interval of 2-3 seconds.

### Data Analysis

All analyses were done using custom MATLAB scripts. Response firing rates were determined by counting the number of spikes over a time window following stimulus onset, converting this count to firing rate (sp/s), and subtracting the baseline firing rate. The window for computing firing rates was visually estimated in order to capture evoked responses for each for each neuron and started, on average, at 120 ms (115 ms) and had a width, on average, of 170 ms (170 ms) for Imc (OTid) neurons. Average firing rates and error bars (s.e.m) were computed from the firing rates across all the repetitions of stimulus presentation, after removing outlier values. Outliers were identified as points that lay outside the range of median ± 1.5*inter-quartile-range of the distribution.

To characterize the responses to the paired presentation of S1 and S2 (i.e., CRPs), we calculated the correlation (Pearson, *corrcoef* command in Matlab) as a function of the strength of S2. A significant negative correlation (p < 0.05) indicated that responses significantly decreased as S2 strength increased. If a neuron did not exhibit significant negative correlation, it was deemed to exhibit fixed response suppression if the suppression was significant for the majority of S2 values (one-way ANOVA on the firing rates to different competitor strengths followed by post-hoc tests against responses to S1 alone, corrected for multiple comparisons). The remaining neurons were considered to not show any effect related to the presence of S2.

Negatively correlated CRPs were fit with a standard sigmoidal function and the parameters of the best fit determined [11]. To obtain reliable estimates of the best fitting sigmoid, any S2 strength for which responses differed non-monotonically from its neighbors, and did so substantially (>150% of the response difference between the neighbors), was omitted from the fitting process and subsequent analyses. We defined transition range of each CRP as the range of strengths of S2 over which responses decreased from 90 to 10% of the max response rate. The half-max of this range was defined as the transition value, i.e., the value of the strength of S2 where responses transitioned from being stronger to weaker than the half-max response. The determination of whether CRPs transitioned abruptly (in a ‘switch-like’ manner) or systematically (in a ‘gradual’ manner) from high to low responses, we adopted previously published conventions [11, 32]: CRPs with transition ranges that were narrower than 4°/s, or 1/5^th^ of the physiological range of S2 strengths were considered to be “switch-like” responses, and others as “gradual” responses. Similarly, for auditory competitors, CRPs with a transition range of 9 dB or less were considered to be switch-like, and the others gradual. Incidentally, these “cut-off” values correspond closely to the dips in the bimodal distributions of transition ranges (Figs. 1F, and 3D, respectively).

For neurons with negatively correlated CRPs, we also calculated the instantaneous firing rates (IFRs) by first obtaining the PSTH of the responses (1 ms time bins), and then smoothing the PSTH with a Gaussian kernel (σ = 12 ms). For each neuron, the IFRs to the paired presentation of S1 and S2 (as well as to S1 alone) were normalized by the peak of the average IFR to S1 alone.

The pooled population responses in Figures 2, 5, and 6 were obtained by binning relative strengths (S2-S1) into five bins, and combining the responses of all the neurons within each bin [11]. This was done separately for OTid and Imc neurons, and separately for neurons with gradual and switch-like CRPs. The time course of response suppression by S2 was determined (within each relative strength bin) by performing a millisecond-by-millisecond ANOVA, comparing the pooled IFRs to S1 alone versus to S1 and S2 presented together [44]. The time-to-suppression was defined as the first millisecond at which the *p-* value of the ANOVA comparison dropped below 0.05, remained below 0.05 for the next 25 ms, and reached 0.01 at least once in that period [11].

Discriminability (d-prime) between responses to two stimulus conditions was computed as the difference in mean responses over the square root of the average of the variances.

For comparison of results from dual, simultaneous recordings in the Imc and OTid, we performed all analyses on data from pairs of multiunit *sites*, rather than on data from pairs of single neurons sorted from these multiunit sites. This is because there was no rational way of establishing which specific neurons sorted from the Imc site ought to be paired with which neurons in the OTid site; comparing all possible pairs would violate the assumption of statistical independence across samples. Notably, because multiunit sites are indeed activated as a whole upon the presentation of stimuli, we do not lose interpretive power in (analyzing and) comparing *site* responses between Imc and OTid.

## ACKNOWLEDGEMENTS

This work was supported by funding from NIH grant R01EY027718. We thank James Garmon for assistance in building the equipment necessary for the experiments and Nagaraj Mahajan for help in data collection and analysis.

